# Accuracy and precision in super-resolution MRI: Enabling spherical tensor diffusion encoding at ultra-high b-values and high resolution

**DOI:** 10.1101/2021.03.17.435819

**Authors:** Geraline Vis, Markus Nilsson, Carl-Fredrik Westin, Filip Szczepankiewicz

## Abstract

Diffusion MRI (dMRI) is a useful probe of tissue microstructure but suffers from low signal-to-noise ratio (SNR) whenever high resolution and/or high diffusion encoding strengths are used. Low SNR leads not only to poor precision but also poor accuracy of the diffusion-weighted signal, as the rectified noise floor gives rise to a positive signal bias. Recently, super-resolution techniques have been proposed for signal acquisition at a low spatial resolution but high SNR, whereafter a higher spatial resolution is recovered by image reconstruction. In this work, we describe a super-resolution reconstruction framework for dMRI and investigate its performance with respect to signal accuracy and precision. Using strictly controlled phantom experiments, we show that the super-resolution approach improves accuracy by facilitating a more beneficial trade-off between spatial resolution and diffusion encoding strength before the noise floor affects the signal. Moreover, precision is shown to have a less straightforward dependency on acquisition, reconstruction, and intrinsic tissue parameters. Indeed, we find that a gain in precision from super-resolution reconstruction (SRR) is substantial only when some spatial resolution is sacrificed. We also demonstrated the value of SRR in the challenging combination of high resolution and spherical b-tensor encoding at ultrahigh b-values—a configuration that produces a unique contrast that emphasizes tissue in which diffusion is restricted in all directions. We conclude that SRR is most valuable in low-SNR conditions, where it can suppress rectified noise floor effects and recover signal with high accuracy. The in vivo application showcases a vastly superior image contrast when using SRR compared to conventional imaging, facilitating investigations of brain tissue that would otherwise have prohibitively low SNR, resolution or required non-conventional MRI hardware.

## Introduction

Diffusion MRI (dMRI) is a non-invasive method for investigating tissue microstructure in healthy and pathological tissue [1]–[3]. Investigations of subtle microstructure features rely on the use of strong diffusion weighting (ultra-high b-values) or tensor-valued diffusion encoding [4]–[6], both of which lead to stronger signal attenuations and low signal-to-noise ratios (SNR). A low signal precision can be improved by averaging over multiple observations which leads to increased scan times. Low SNR also causes poor signal accuracy in magnitude imaging due to the so-called rectified noise floor, which induces a positive signal bias [7]. Unlike precision, signal accuracy is not improved by averaging over magnitude signals [8]. Although averaging over complex signals with a coherent phase would improve accuracy, it is challenging because the diffusion encoding causes phase variation in the presence of tissue motion [9][10]. Accurate measurements of the diffusion-encoded signal is thus challenging.

One approach where the problem of low SNR is particularly acute is spherical b-tensor encoding at ultrahigh b-values. This combination is desirable because it provides a novel contrast that emphasizes tissue in which diffusion is restricted in all directions. For example, it can be used to highlight the tightly packed granule cells in the cerebellar cortex, which are affected in diseases such as spinocerebellar ataxis and Alzheimer disease [11], [12]. So far, this contrast has been obtained only at preclinical MRI systems [13], or systems with ultra-strong gradients and at a poor spatial resolution [6]. Making this contrast available at high resolution and at widely available clinical MRI systems would add a new tool to the neuroimaging toolbox and enable studies of the cerebellum in a wide range of neurological conditions.

Super-resolution reconstruction (SRR) is a promising solution to the problem of low SNR in dMRI. In principle, SRR is based on data acquired at a low spatial resolution—with improved precision and accuracy—and a subsequent image reconstruction that recovers a high-resolution image. SRR methods can balance the trade-off between SNR, spatial resolution and acquisition time [14], [15], [24], [16]–[23][25]. Wu et al. [21] used high-order singular value decomposition to regularize a patch-based SRR framework, while Yang et al. [22] proposed a non-local strategy where joint information from the adjacent scanning directions was used to improve resolution. Poot et al. [24] demonstrated increased resolution of diffusion tensor parameters from a set of super-resolved diffusion-weighted images, where each image is reconstructed from set of low-resolution images with the same diffusion weighting and gradient direction. Van Steenkiste et al. [14] showed increased spatial resolution of diffusion tensor parameters when optimizing both k- and q-space sampling, and Jeurissen et al. [15] showed improved accuracy and precision in q-space trajectory imaging parameter estimation. Looking at this research, we note that the acquisition of the low-resolution data can be designed in several ways. For example, the image acquisition can be performed with three slices having orthogonal low-resolution axes [25], slice shifting along the low-resolution direction [18], or multiple stacks of slices rotated about a common axis [16]. This flexibility of SRR allows it to be tailored to the needs of echo-planar imaging (EPI) which is commonly used in dMRI. For example, it is convenient acquire data using a common phase encoding direction to avoid that local field inhomogeneities cause variable geometric distortions [26].

Although SRR has seen a broad uptake, the literature lacks a systematic treatment of SRR noise propagation. For example, imaging at a low resolution allows faster imaging (shorter repetition times), but leads to a complex interplay between SNR, scan time, repetition time, and T1-relaxation. A careful investigation of this interplay is necessary to understand the acquisition trade-offs, and to leverage them for optimal experimental design. Furthermore, the application of SRR for spherical b-tensor encoding at ultrahigh b-values and high resolution are yet to be explored. In this work, we aim to describe a general framework for SRR, formally analyze noise propagation, and experimentally verify the impact of SRR on signal accuracy and precision. We demonstrate the value of SRR in the challenging combination of spherical b-tensor encoding at a b-value of 4.0 ms/μm^2^ and 1.6 mm^3^ isotropic resolution in the brain.

## Theory

Super resolution reconstruction aims at recovering a high resolution image from multiple low-resolution images that sample the object in different ways. A common approach is to acquire multiple stacks of thick slices rotated around the phase encoding direction [16]. To reconstruct an image with isotropic voxel sizes, the lower limit of low-resolution rotations *N_R_* is given by [19]

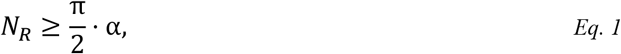

where α is the aspect factor, defined as the ratio between the voxel size in the slice direction and frequency/phase encoding direction. Note that this factor also captures the volume ratio between the low and high-resolution voxels, as in-plane resolutions are identical. Also note that SNR is proportional to voxel volume in multislice acquisitions [27].

The mapping from a high resolution image (**x**) to a low resolution image (**y**) can be described by a linear system [28]

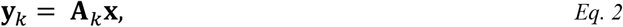

where *k* is an index of the low-resolution image sample. The sampling matrix **A**_*k*_ describes the rotation/translation, down-sampling and blurring of the underlying high-resolution object and can be constructed from the pulse sequence settings. Both **y**_*k*_ and **x** are expressed as column vectors such that a square image with *N* voxels on each side is represented by a *N*^2^×1 vector. The complete sampling of all low resolution images can be described in a single linear system

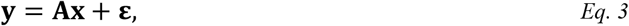

where **ε** is random noise. Assuming the noise is independent and normally distributed with zero mean, the solution to recover **x** given **A** and ***y*** can be expressed as a least squares problem, such that

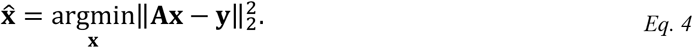

The greater the number of complementary observations in **y**, the better the condition of the problem becomes. However, the problem in Eq. 4 often remains ill-posed due to the down-sampling operation included in **A**. Therefore, the solution requires regularization, which often translates to imposing a smoothness to the solution [20]. A common approach is Tikhonov regularization [29], which penalizes high spatial-frequencies in the estimated high-resolution image. Including this regularization, the regularized least-squares squares problem becomes

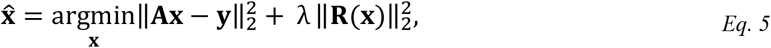

where **R** is the regularization term and λ is a scalar weight. We will here use a general regularization term independent of the image content: **R(x)** = **I**, where **I** is the identity matrix. This enables the use of the closed form solution, according to

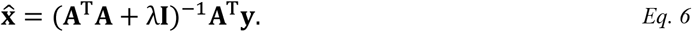

Note that **A^T^y** produces the average low-resolution signal on the high-resolution grid, i.e., a blurred image. Without regularization (λ = 0), the remaining term (**A^T^A**)^-1^ is a sharpening operator, ideally reproducing the true image when applied to **A^T^y**. As λ increases, the sharpening is reduced. However, as λ alters the denominator in Eq. 6, the intensities in 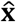 are dependent on λ. To remove this dependence and thereby simplify comparisons among sampling schemes as described later, we can rewrite Eq. 6 according to

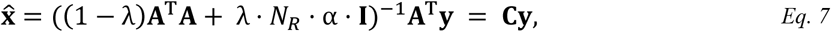

where **C** contains the entire reconstruction operation. The new regularization factor is constrained to 0 ≤ λ ≤ 1, such that λ = 1 merely returns the average low-resolution signal on the high-resolution grid, but corrected for the intensity gain caused by larger voxel volumes. Figure 1 illustrates the SRR process for an in vivo acquisition for weak, moderate, and strong levels of regularization (different values of λ). Weak regularization amplifies noise while strong regularization results in a blurred image. Moderate regularization balances the two.

**Figure 1.**
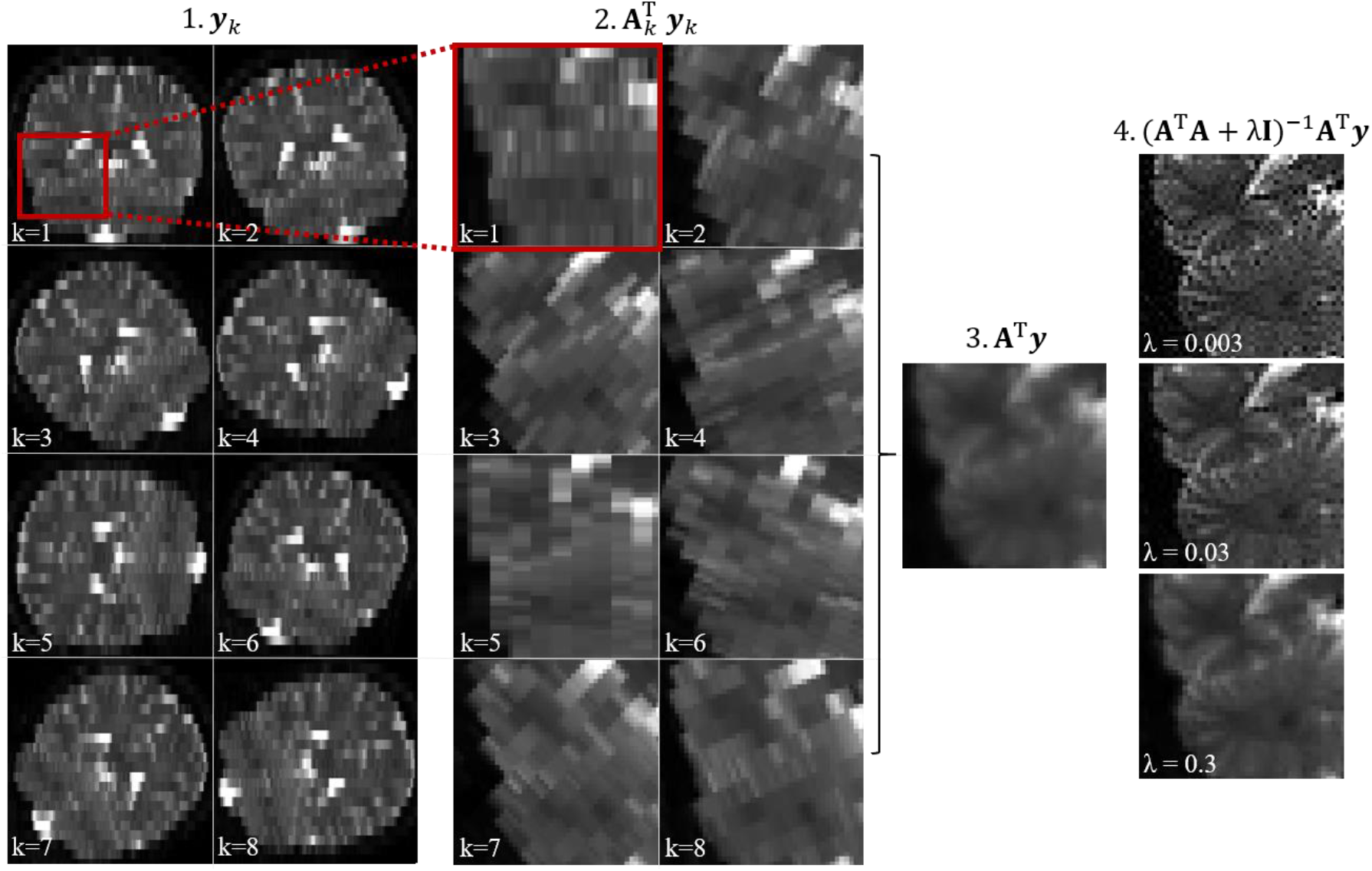
Illustration of the super-resolution reconstruction process. Step 1: Multiple images are acquired with low through-plane resolution rotated around a static axis. Step 2: The images are up-sampled to the high-resolution grid by application of the upsampling operator to each individual image. Step 3: The joint up-sampling operator results in an average of the individual images, i.e., a smooth image on the high-resolution grid. Step 4: The sharpening operator is applied to obtain a high-resolution image. The regularization parameter λ determines the trade-off between resolution and noise propagation; a higher λ leads to a blurrier, but less noisy image. Steps 2 to 4 are shown as a magnified view of the region indicated by the red square.

The effect of regularization on the effective spatial resolution of 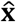 can be characterized using impulse response analysis, in which an impulse signal is passed through the forward model **A** and reconstruction matrix **C** according to

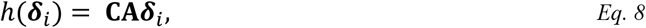

where ***δ**_i_* is an impulse vector at position *i* of **x** and *h* describes the resulting point spread function. Ideally, this would return the same ***δ**_i_*, meaning no loss of resolution due to reconstruction. Note that the point spread function is dependent on location, properties of the sampling (**A**, *N_R_*, α), and the level of regularization (λ).

### Signal accuracy and precision

We evaluate the performance of SRR in terms of signal accuracy and signal precision. Across repeated measurements under identical conditions, signal *accuracy* (or trueness)^1^ concerns the closeness of the average signal to the true value, while signal *precision* concerns the spread of the signal. Both terms are strongly influenced by the data distribution that characterizes the MR signal. The noise in complex MR signal is normally distributed, whereas the magnitude signal used in dMRI is approximately Rice distributed [7] (for a detailed review of MR data distributions, see [30]). Consequently, in the absence of true signal, the mean (η) magnitude signal is given by [31]

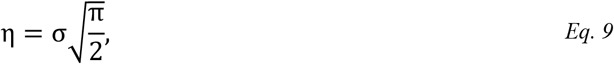

where σ > 0 is the standard deviation of the measured signal. η is commonly referred to as the rectified noise floor, which gives rise to a positive signal bias that becomes prominent at low SNR. In presence of true signal, an approximation of the mean of the measured signal 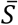 is given by [7]

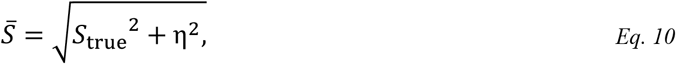

where *S*_true_ is the signal in the absence of noise. Hence, as a measure of signal accuracy, we employ the signal-to-noise-floor ratio (SNFR) [32]

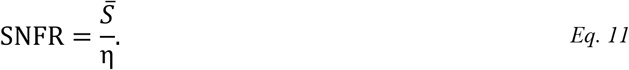

Note that the SNFR of a non-diffusion weighted measurement characterizes the maximum achievable signal attenuation for accurate signal sampling. As a measure of precision, we use the signal-to-noise ratio (SNR) defined as

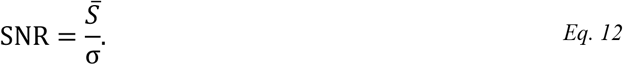

Note that SNR and SNFR are both proportional to the voxel volume whereas noise levels are independent of the voxel volume [27].

To enable comparison of precision between SRR and a conventional high-resolution acquisition (referred to as direct sampling), we define the SNR efficiency factor (ρ), according to

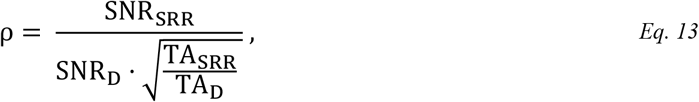

where SNR_SRR_ and SNR_D_ are the SNR levels and TA_SRR_ and TA_D_ the acquisition times of SRR and direct sampling at a given spatial resolution. To evaluate Eq. 13, we study how noise propagates from **y** into **x**. For SNR > 3 the noise distribution is approximately Gaussian, independent and identically distributed [7][33]. The signal variance in the reconstructed image can thus be easily computed from the linear operations in Eq. 6 based on the additive property of variance. We define the noise propagation factor (*κ*) as the average ratio of the noise standard deviation in the high-resolution reconstructed and low-resolution images, according to

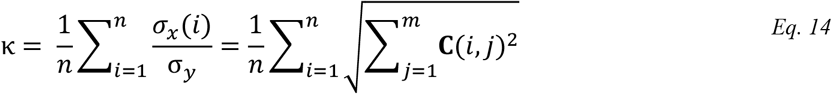

where *n* is the total number of reconstructed voxels with index *i*, and *m* is the total number of low-resolution input voxels with index *j*. The noise propagation factor is, like the point spread function, dependent on properties of the sampling (**A**, N_R_, α) and the regularization (λ). For a setup where the only difference between direct sampling and SRR is related to the SRR configuration (slice thickness, number of slices, repetition time and number of slice orientations), Eq. 13 can be extended to include the relevant effects of imaging parameters (Appendix A) to

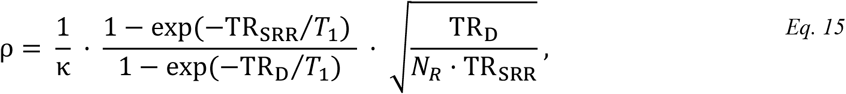

where TR_D_ and TR_SRR_ are the repetition times for direct sampling and SRR respectively, and *T*_1_ is the longitudinal relaxation time. The TR is approximately proportional to the number of slices, and the number of slices to cover the same area in an SRR acquisition can be reduced with a factor of α. Therefore, the minimal TR_SRR_ is given by

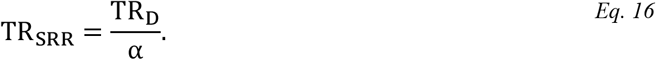

When TR_D_ » *T*1, both 1 — exp(—TR_SRR_/*T*_1_) and 1 — exp(—TR_D_/*T*_1_) will approach unity, as will their ratio. When sampling directly, approximately 1% signal is lost due to incomplete T1 recovery when TR_D_/*T*_1_ = 4.6. However, this corresponds to a signal loss of 46% when using α = 6 and a minimal TR_SRR_. For this reason, as the aspect factor increases and the TR is minimized, the expected SRR gain will eventually be negated by incomplete T1-recovery.

## Methods

We evaluate the performance of the outlined SRR framework. First, we investigate signal accuracy in a water phantom, both numerically and experimentally. Second, we investigate signal precision in terms of SNR efficiency, both numerically, analytically, and experimentally in vivo. Finally, we demonstrate the utility of SRR at spherical tensor dMRI at ultrahigh b-values in vivo. All simulations and data analysis were performed in Matlab (The MathWorks, Inc., Natick, Massachusetts, USA).

### Data acquisition

All practical experiments were performed on a 3T-scanner (MAGNETOM Prisma, Siemens Healthcare, Erlangen, Germany) using a 20-channel head and neck coil. The study was approved by the local ethics committee and informed consent was obtained from all volunteers. A prototype pulse sequence was used based on a single-shot spin-echo with echoplanar imaging readout that facilitates user defined gradient waveforms for diffusion encoding [34]. Gradient waveforms for spherical b-tensor encoding were optimized for the MRI system [35], including compensation for concomitant gradients [36]. The waveforms were constrained to a maximal gradient magnitude of 80 mT/m (L2-norm) and a slew rate of 100 T/m/s. The gradient waveform is shown in Appendix B. Detailed information on acquisition parameters will be described per experiment.

Low-resolution data was acquired with slices rotating around a fixed phase encoding direction. As experiments were performed for various aspect factors, we used the minimum number of low-resolution rotations per aspect factor according to Eq. 1. In simulations, Eq. 2 was used to obtain low-resolution data. In all experiments, low-resolution data was reconstructed per slice according to Eq. 7.

### The impact of noise floor on signal accuracy

We characterize the effect of the rectified noise floor on signal accuracy when modulating the aspect factor as well as the strength of diffusion encoding. We use a mono-exponential signal model in water given by

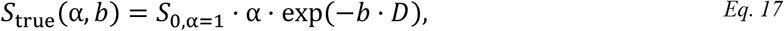

where *b* is the b-value and *D* is the mean diffusivity. We include Rice noise contribution to obtain a simplified analytical signal model to fit measured data, according to Eq. 10.

Measurements were performed in a phantom filled with deionized water at room temperature. Data were acquired in a single slice with an in-plane resolution of 1.6×1.6mm^2^ and slice thicknesses between 1.6 mm and 9.6 mm, giving aspect factors between 1 and 6. A constant TR of 4 s was used, to remove influence of T1-relaxation. A single slice acquisition was used to ensure that all aspect factors shared the same central plane and to provide a fair comparison of baseline signals. We used b-values ranging from 0 to 3 ms/μm^2^ in steps of 0.3 ms/μm^2^ and 5 repetitions. The average signal *S*(α, *b*) was estimated separately for each b-value in a homogeneous area of the phantom. The diffusivity *D* and noise level *σ* were estimated by a least-squares fit of the data to Eq. 10 given Eq. 17.

The same conditions were reproduced in a numerical signal model where *S*_0,α=1_ = 1 and *D* = 2.2 μm^2^/ms. Noise with *σ* = 0.014 was added to the real and imaginary channel for signal generated by Eq. 17, after which the magnitude was computed. This corresponded to a maximal SNR of *S*_0_/*σ* = 71 and 426 for α = 1 and 6 respectively. The average signal *S*(α, *b*) was estimated from 10^4^ realizations of noise. Note that diffusivity and noise levels were matched to those measured in the water phantom.

The numerical signal model was extended to include SRR. We simulated low-resolution measurements in a Shepp-Logan phantom, where one of the regions was adapted to mimic water at room temperature. High-resolution images were reconstructed for 10^3^ realizations of noise, after which the average signal *S*(α, *b*) was estimated.

From all resulting signal curves, we estimated the SNFR (Eq. 11) and computed the threshold attenuation factor (b_max_ · D) at which the signal bias is less than 5% using the true mono-exponential signal model as reference. The threshold attenuation factors were compared across aspect factors.

### Analysis of precision and SNR efficiency

We investigated SNR efficiency (Eq. 13–Eq. 15) under assumption of Gaussian noise (SNR > 3). To avoid unfair gains in precision due to the regularization in SRR, we compared SNR efficiency at matched effective spatial resolutions. We used the full-width-half maximum (FWHM) of the average point spread function (Eq. 8) in the anterior-posterior direction as a measure of spatial resolution. To match resolutions between SRR protocols, we searched for the λ that induced a FWHM matching the reference value. For direct sampling, we matched resolution by convolving the images by a gaussian filter with a size dictated by the FWHM. In simulations, we ensured matched resolutions by convolving the directly sampled data with the average point spread function per SRR protocol.

We performed multiple experiments to investigate the relationship between SNR efficiency, aspect factor, T1-relaxation and acquisition time. When T1-relaxation is neglected, we expect that SNR efficiency increase with aspect factors monotonically. Simulations were performed in a Shepp-Logan phantom, where low-resolution data was sampled for aspect factors between 1 and 8 assuming *S* ∝ α and *S*_α=1_ = 1. Gaussian noise with *σ* = 0.1 was added and the SNR efficiency was estimated from the reconstructed images in a homogenous area according to Eq. 13, where we set 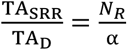. The procedure was repeated for weak, moderate, and strong regularization, associated with an average point spread function with a FWHM of approximately 1.17, 1.35 and 1.59 voxels, respectively. The estimation of SNR efficiency was also performed analytically using Eq. 15. In the same analytical model, we also included T1 relaxation effects, where we expect that smaller TR reduces SNR efficiency. We evaluated SNR efficiency at moderate regularization for TR_D_ = [5 10 20] s and TR_SRR_ set to the minimum possible value for each aspect factor (Eq. 16). T1 was set to 1.6 s, as observed in brain grey matter at 3 T [37]. Since TR_SRR_ does not have to assume the minimal value, we also investigated how longer TR_SRR_ promotes T1 relaxation and how it affects the SNR efficiency in both grey and white matter (T1_WM_ = 0.8 s [37]) for α = 8.

To verify our simulations, SNR efficiency was also evaluated in a healthy brain in vivo (male, 28 years) for aspect factors up to 6. All experiments used *b* = [0 0.5] ms/μm^2^ with 1 and 10 repetitions, FOV = 220×220×144 mm^3^, TE = 100 ms, partial-Fourier factor = 6/8, 2x in-plane acceleration (GRAPPA) and bandwidth = 1725 Hz/pixel. Remaining imaging parameters dependent on the aspect factor are summarized in Table 1. All images were reconstructed at a resolution of 1.6×1.6×1.6 mm^3^ to yield a point spread function with an FWHM of approximately 2.2 mm due to regularization. The SNR efficiency was estimated according to Eq. 13 in the central white matter, brainstem, corpus callosum and cerebellar white matter at b = 0.5 ms/μm^2^.

**Table 1.** IN VIVO ACQUISITION AND RECONSTRUCTION PARAMETERS FOR SUPER-RESOLUTION RECONSTRUCTION PROTOCOLS

| | $\alpha=1$ | $\alpha=2$ | $\alpha=3$ | $\alpha=4$ | $\alpha=5$ | $\alpha=6$ |
| --- | --- | --- | --- | --- | --- | --- |
| $N_R$ [1] | 1 | 3 | 5 | 7 | 8 | 10 |
| Res [mm<sup>3</sup>] | 1.6×1.6×1.6 | 1.6×1.6×3.2 | 1.6×1.6×4.8 | 1.6×1.6×6.4 | 1.6×1.6×8.0 | 1.6×1.6×9.6 |
| TR [ms] | 13000 | 6500 | 4300 | 3300 | 2600 | 2200 |
| TA [min] | 2:23 | 3:35 | 4:00 | 4:14 | 3:49 | 4:02 |
| $\lambda$ [1] | - | 0.19 | 0.10 | 0.06 | 0.05 | 0.04 |

### In vivo dot fraction imaging

We demonstrate the utility of SRR by visualizing the presence of the so-called dot fraction in the brain with a higher contrast than possible when using direct sampling. The dot-fraction is linked to the relative signal that remains at very high b-values when using spherical b-tensor encoding. As anisotropic tissue is effectively attenuated by spherical b-tensor encoding, the remaining signal can be attributed to restricted pools in which the apparent diffusivity is low or zero in all directions [6] [50].

A healthy volunteer (male, 27 years) was scanned at b = [0 1 4] ms/μm^2^ using 1, 3, and 13 repetitions, *N_R_* = 8, resolution = 1.6×1.6×7.2 mm^3^ (α = 4.5), FOV = 211×211×144 mm^3^, TE = 120 ms, TR = 4200 ms, TA = 9:31 min, 2x in-plane acceleration (GRAPPA), partial-Fourier = 6/8 and bandwidth = 1720 Hz/pixel. A directly sampled set was acquired for comparison at b = [0 1 4] ms/μm^2^ using 1,4, and 15 repetitions, resolution = 1.6×1.6×1.6 mm^3^, FOV = 211×211×188 mm^3^, TR = 14200 ms and TA = 9:18 min. The TR was set to the minimum value possible without the use of through-plane acceleration. To reduce the impact of system drift, we used 2x through plane acceleration and interleaved the b-values over volumes [38][39], [40]. All raw data were denoised using Marchenko-Pastur principle component analysis [41], [42]. The low-resolution images were reconstructed at a resolution of 1.6×1.6×1.6 mm^3^ using SRR, where we set λ = 0.05 (FWHM of point spread function is 2.1 mm).

We assume a dot compartment with signal fraction *f*_dot_ and isotropic diffusivity *D*_dot_ equal to zero, accompanied by a fraction of other tissue (1 — *f*_dot_) with non-zero isotropic diffusivity. Assuming Gaussian diffusion and no exchange, the diffusion-weighted signal *S*(*b*) is given by [6], [43]

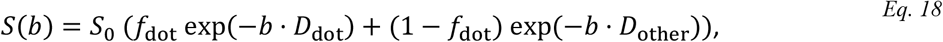

where *S*_0_ is the non-diffusion weighted signal. For very high b-values, the exp(—*b*_high_ · *D*_other_) ≈ 0, and only signal in the dot compartment remains since *D*_dot_ ≈ 0, which simplifies Eq. 18 to

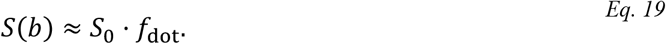

The resulting map is weighted by both *S*_0_ (T2-weighting) and *f*_dot_ (influenced by density of cells exhibiting restricted diffusion in all directions). Note that, under these assumptions, *f*_dot_ only gives an upper limit on the true value as other tissues as well as the rectified noise floor may contribute to the remaining signal [6].

Both the super-resolved and directly acquired high-resolution images were averaged for *b* = 4 ms/μm^2^. The grey-to-white matter signal ratio was calculated between gray matter voxels in the cerebellar cortex and white matter voxels in the cerebellum. *f*_dot_ was calculated using Eq. 19. We compared the images to a corresponding T1-weighted morphological scan and to a similar contrast found in brain histology from the BigBrain atlas [44], Nissl stained to emphasize neurons.

## Results

### The impact of noise floor on signal accuracy

Figure 2 shows the effect of the rectified noise floor on signal accuracy for different aspect factors. The rectified noise floor causes an overestimation of the signal at high b-values where SNR is low (Fig. 2a). As the SNFR is boosted by a factor α, higher b-values can be employed before reaching the 5% signal bias threshold: for the measured signal (SNFR = 60 at *b* = 0 for α = 1, *D* = 2.2 μm^2^/ms), sampling with α = 6 compared to α = 1 allows for a b-value increase from 1.4 to 2.2 ms/μm^2^ (Fig. 2b). More generally, going from α = 1 to α = 6 allows for an increase of the attenuation factor 80%. Simulated results agree with measurements. Note that TR was constant across measurements and does not reflect the contribution from T1-weighting.

**Figure 2.**
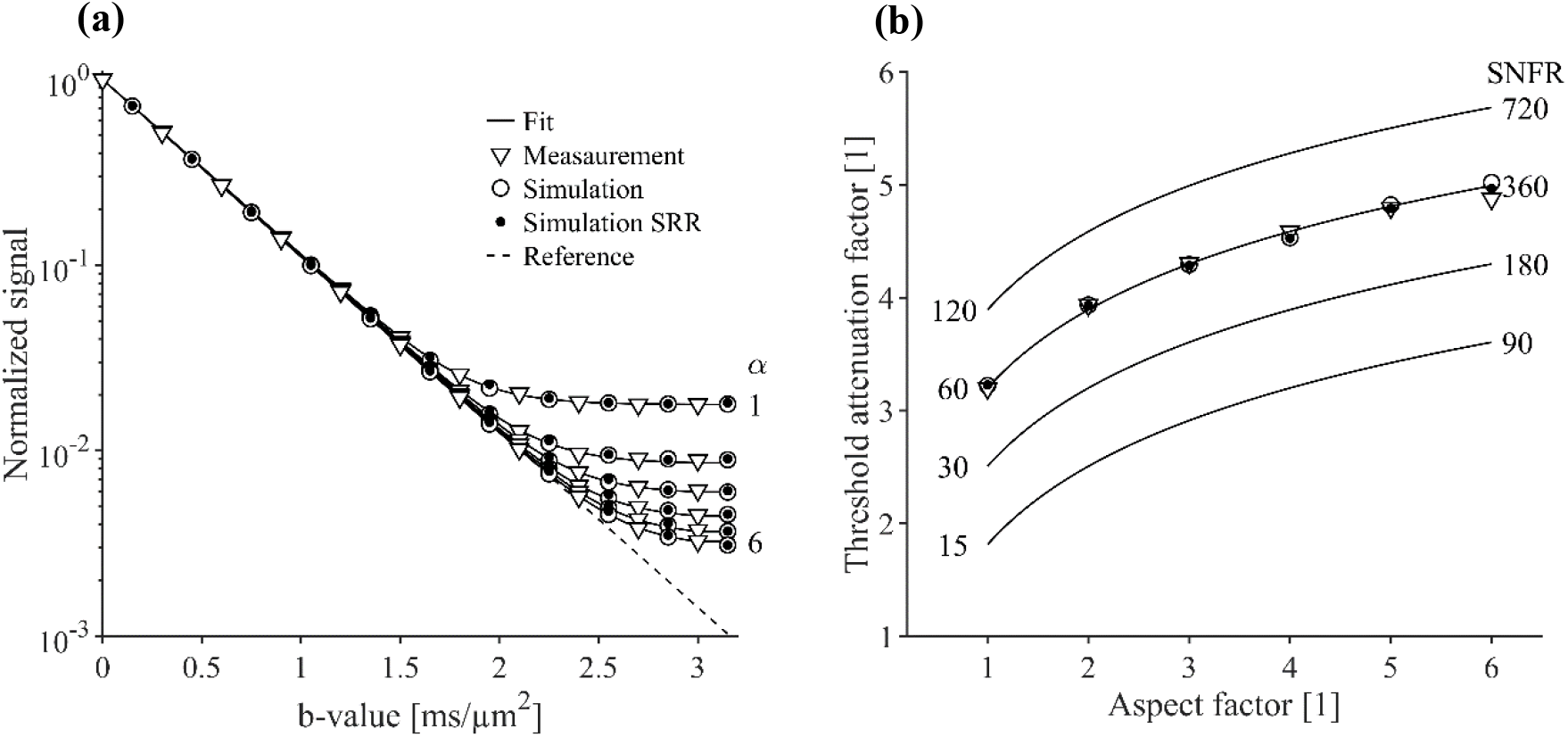
Effect of the rectified noise floor on signal accuracy for aspect factors (α) up to 6. Panel (a) shows the simulated and measured mean signal in a water phantom as a function of the b-value. Panel (b) shows the threshold attenuation factor (*b_max_* · *D*) that can be used for accurate signal sampling for different SNFR at b0 versus aspect factor. Sampling with larger voxels improves accuracy and allows for the use of higher b-values before the noise floor affects the signal. Simulations and experimental results (circles versus triangles) show high agreement.

### Analysis of precision and SNR efficiency

Figure 3 shows the SNR efficiency for SRR with different aspect factors for three levels of regularization. SNR efficiency generally increases with aspect factor. For example, for moderate regularization (FWHM of the point spread function is 1.35 voxels), ρ ≈ 2 for α = 8, effectively doubling the precision. As expected, a stronger regularization leads to higher SNR efficiency but lower effective spatial resolution. Note that this increase is not just due to the smoothing induced by strong SRR regularization. The comparison was made at matched effective resolutions, which shows that regularized SRR is more SNR effective than smoothing a direct acquisition. However, this analysis does not include T1-relaxation effects.

**Figure 3.**
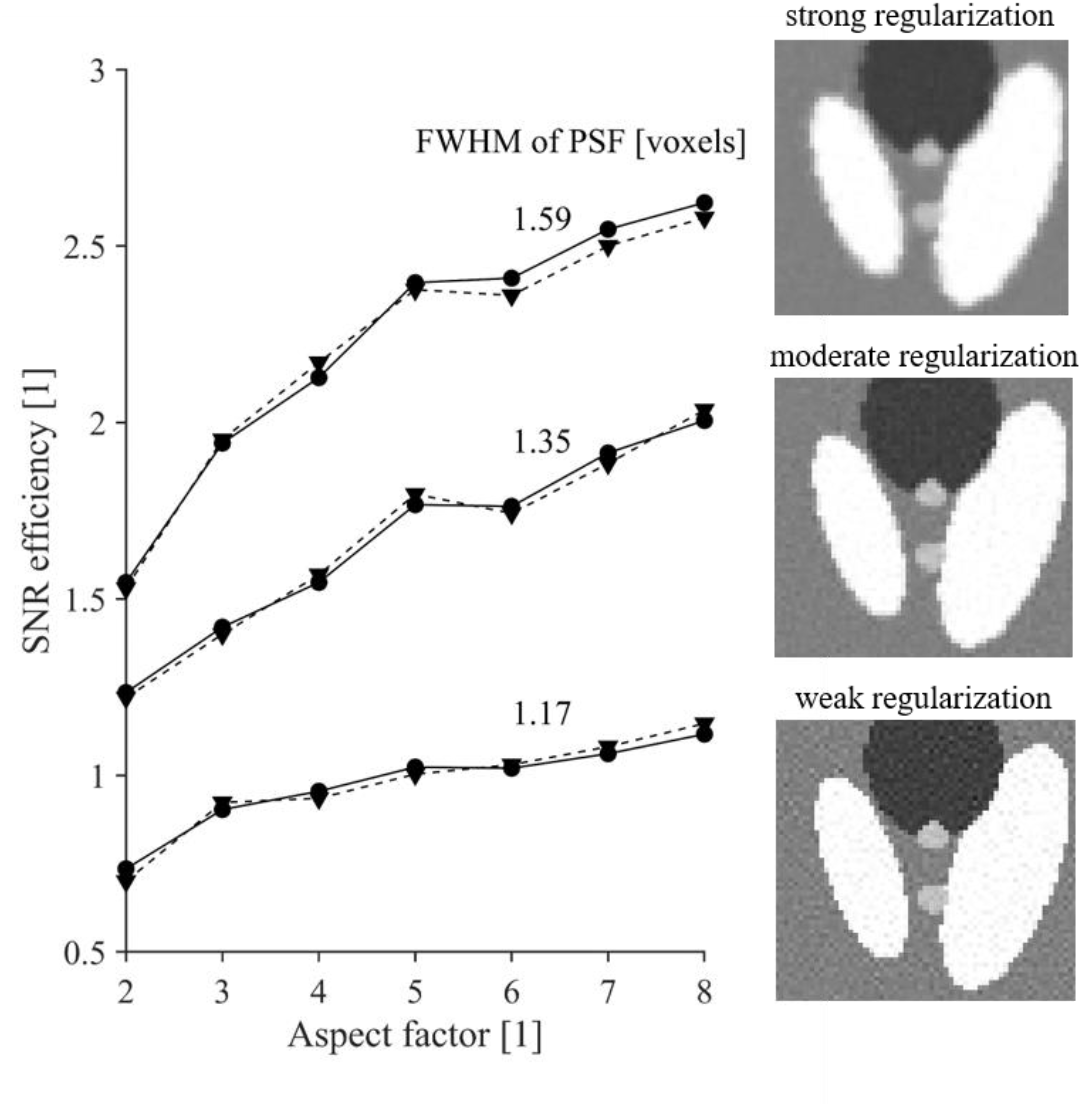
SNR efficiency for different aspect factors and regularization strengths, evaluated numerically (dashed line) and analytically (solid line). SNR efficiency above unity means that precision increases compared to a direct acquisition with matched spatial resolution and acquisition time. Both the increase in aspect factor and regularization strength lead to a higher SNR efficiency. Analytical and numerical experiments agree. Effects of T1-relaxation are disregarded in this analysis.

Figure 4 shows the effect of T1 saturation on SNR efficiency, which decrease with TR_D_ (Fig. 4a). This effect is more evident for higher aspect factors. For example, for α = 8 in grey matter, ρ = 2 at TR_D_>>T1 decreases to ρ = 0.7 at TR_D_ = 5 s. Rather than increasing SNR efficiency, SRR in this case leads to a reduced SNR efficiency. However, when TR_D_ > 10 s and/or lower aspect factors are used, T1 effects are less evident and SNR efficiency is still increased for SRR. Note that TR_SRR_ can be increased at the expense of scan time. As illustrated for α = 8, we see it is only beneficial to set TR_SRR_ to a minimum when TR_SRR_/T1 > 1.25 (Fig. 4b).

**Figure 4.**
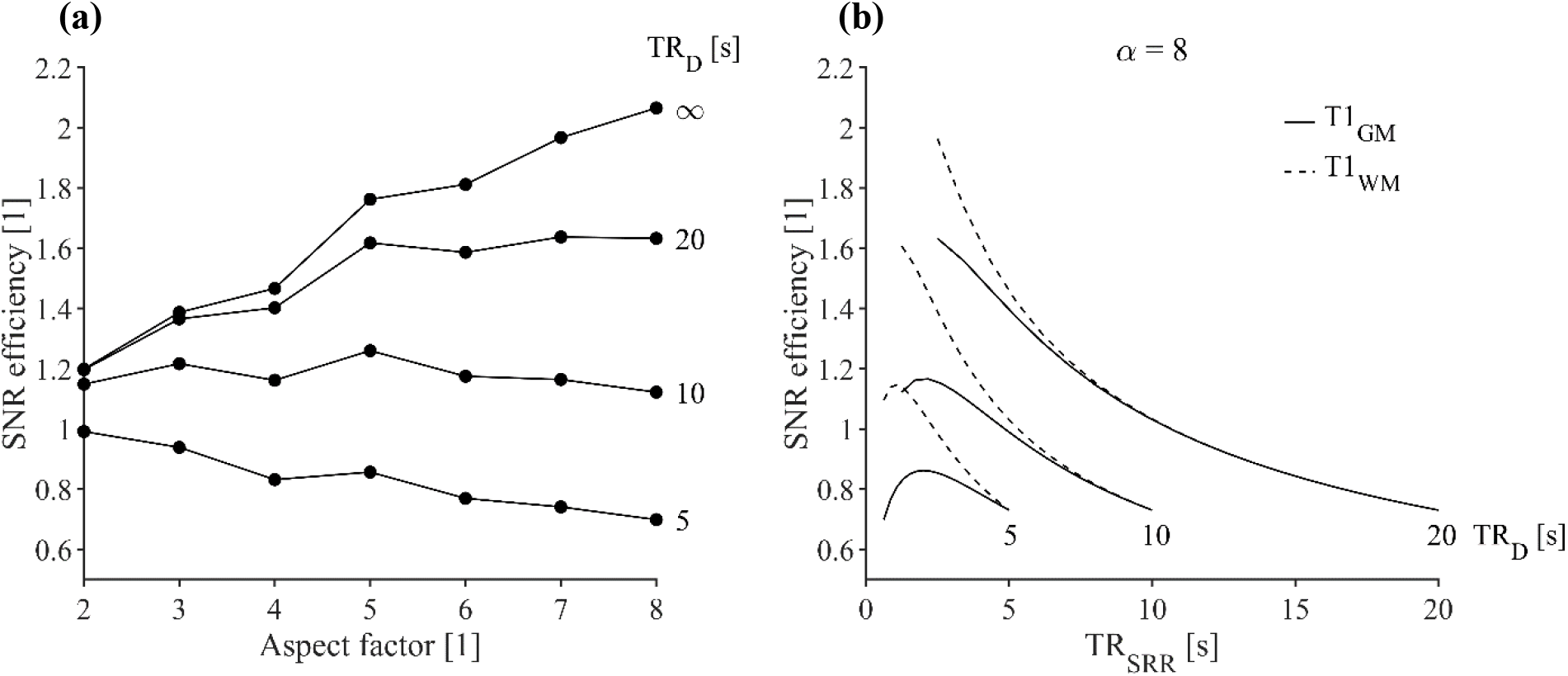
Effect of T1 relaxation on the SNR efficiency. Panel (a) shows the SNR efficiency in grey matter using the minimum TR_SRR_ for a given TR_D_. and aspect factor. SNR efficiency decreases as TR_D_ decreases, with a faster decrease for higher α. Panel (b) shows the effect of changing TR_SRR_ above its minimum for α = 8 and T1 of both grey- and white matter. SNR efficiency is maximized when TR_SRR_/T1 is as close as possible to 1.25. Simulations are performed at moderate regularization strength (FWHM of PSF is 1.35 voxels).

### In vivo results on precision

Figure 5 shows the results of SRR in vivo for different aspect factors but similar effective resolution (FWHM matched). It shows that SRR in vivo is feasible, as resolution is regained for all aspect factors. Ringing artefacts are observed near high-contrast transitions, such as around the ventricles [45]. A variable T1-weighting can be seen as the aspect factor increases and the TR_SRR_ is reduced.

**Figure 5.**
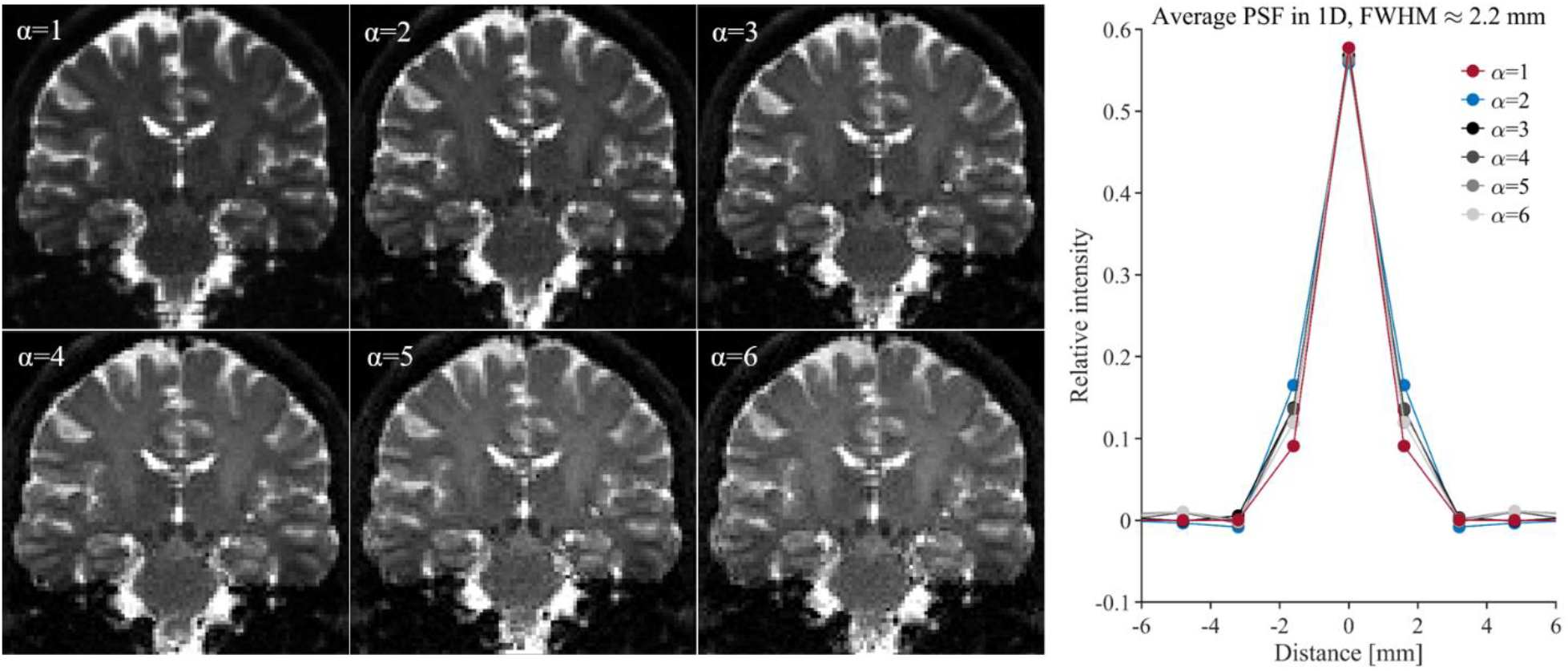
Super-resolution reconstruction (SRR) in vivo at *b* = 0 ms/μm^2^. Panel (a) shows images acquired without diffusion weighting and reconstructed at 1.6×1.6×1.6 mm^3^ using different aspect factors (α). Contrast reduces as the aspect factor grows, due to T1-relaxation effects. Panel (b) shows the average point spread functions induced by the different SRR protocols. The PSFs are aligned between protocols, i.e. the images are given at an equal resolution. Note that α=1 refers to a Gaussian smoothened direct acquisition.

Figure 6 shows SNR efficiency for measured and simulated data in white matter at *b* = 0.5 ms/μm^2^. In central white matter and the corpus callosum, experiments and simulations agree up to α = 5. In the brainstem and cerebellar white matter, simulations overestimate the SNR efficiency for all aspect factors. This could partly be explained by differences in the T1-times of the underlying tissue. For example, the T1 in brainstem is reported to be 1.2 s [46] compared to a T1 of 0.8 s in central white matter used for simulations, thereby reducing SNR efficiency according to Eq. 15. In all cases, the SNR efficiency is still above unity, meaning SRR is beneficial over direct sampling for this specific protocol.

**Figure 6.**
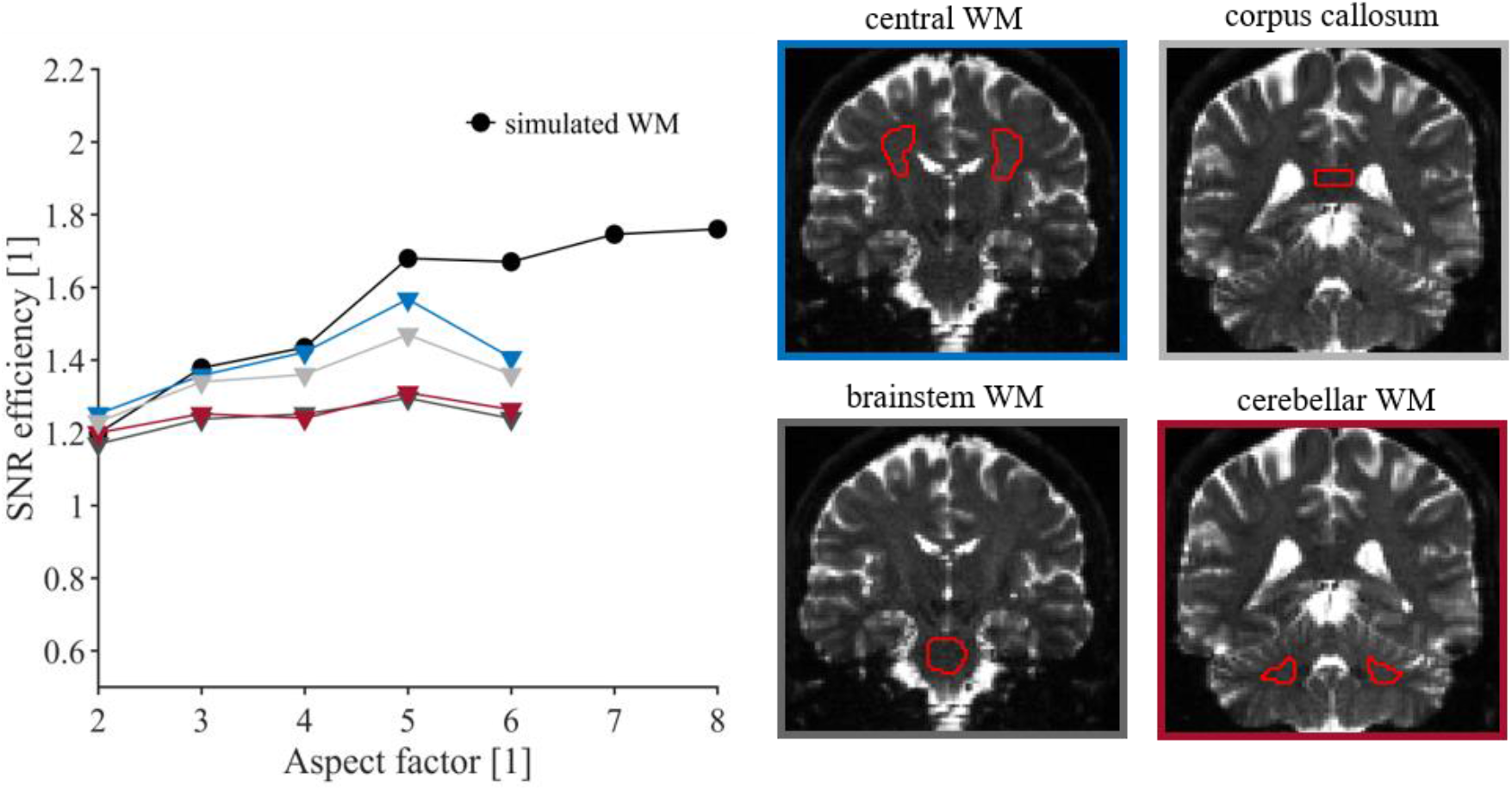
SNR efficiency of white matter (WM) in vivo. The values were obtained from simulations as well as estimated from experimental data, for different aspect factors. Generally, simulations overestimate the SNR efficiency, which could be due to T1 differences of the underlying tissue. Note that in all cases SNR efficiency is above unity, meaning precision is gained by SRR compared to direct sampling.

### In vivo dot fraction imaging

Figure 7 shows the results for dot fraction imaging. Direct sampling leads to poor image contrast throughout the brain. By contrast, SRR enables a vastly improved image contrast where the cerebrum becomes visible with prominent signal in the cerebellar cortex. The contrast ratios between the cortex and white matter of the cerebellum are 1.82 for SRR and 1.06 for direct high-resolution. Figure 8 shows that a part of this contrast is due to T2 effects, as the T2-adjusted map of *f*_dot_ shows a less pronounced contrast. A similar contrast is observed in neuron-stained brain histology.

**Figure 7.**
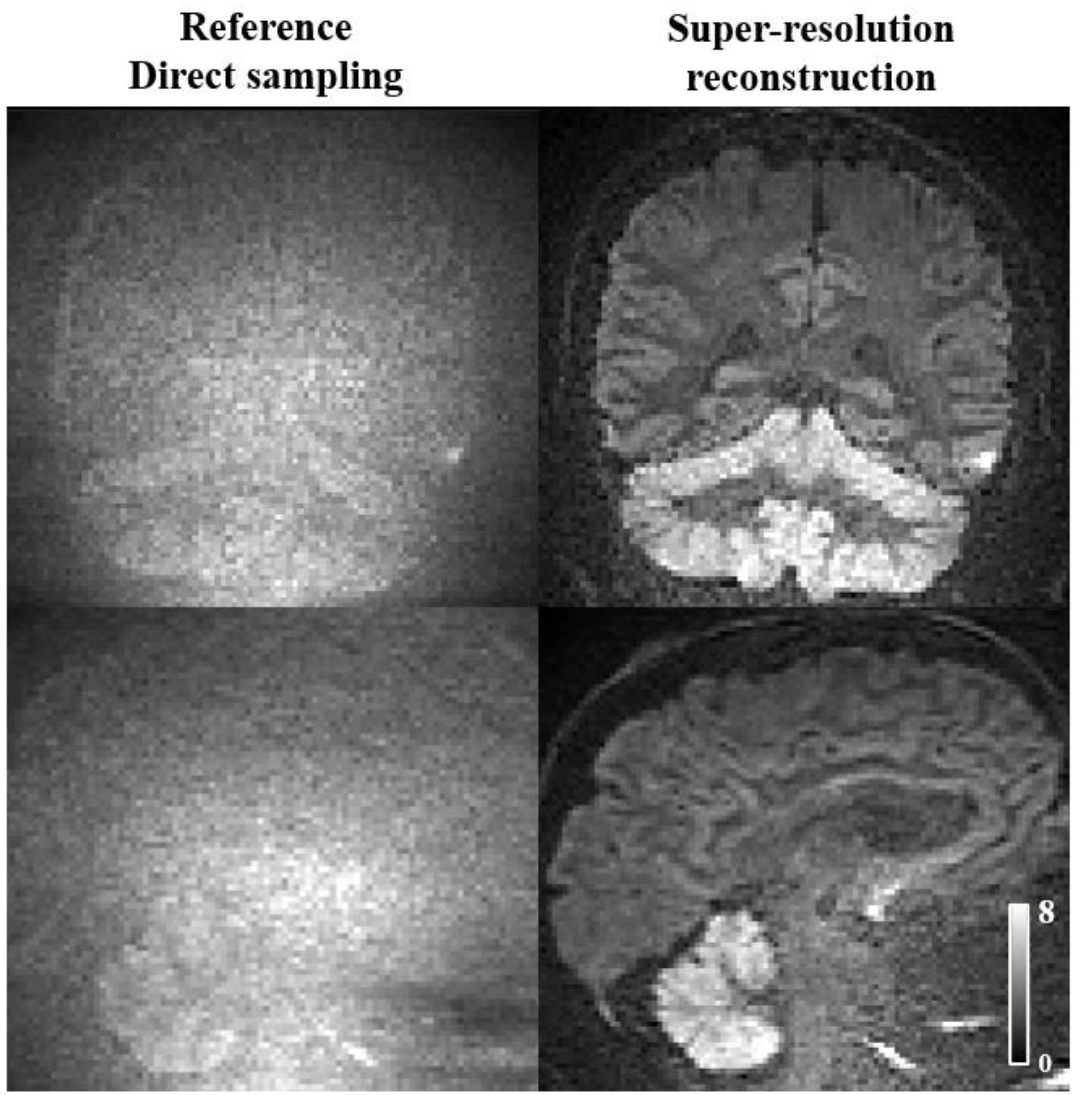
In vivo illustration of the benefits of SRR. The figure shows diffusion weighted images from a direct acquisition (left) and an SRR protocol (right) with spherical encoding at *b* = 4 ms/μm^2^ in coronal and sagittal view at 1.6×1.6×1.6 mm^3^. A vastly higher contrast is observed with SRR compared with direct sampling. Quantitatively, this corresponds to an increase in the contrast ratio between the cerebellar cortex and white matter to 1.82 from 1.06. Acquisition times are similar.

**Figure 8.**
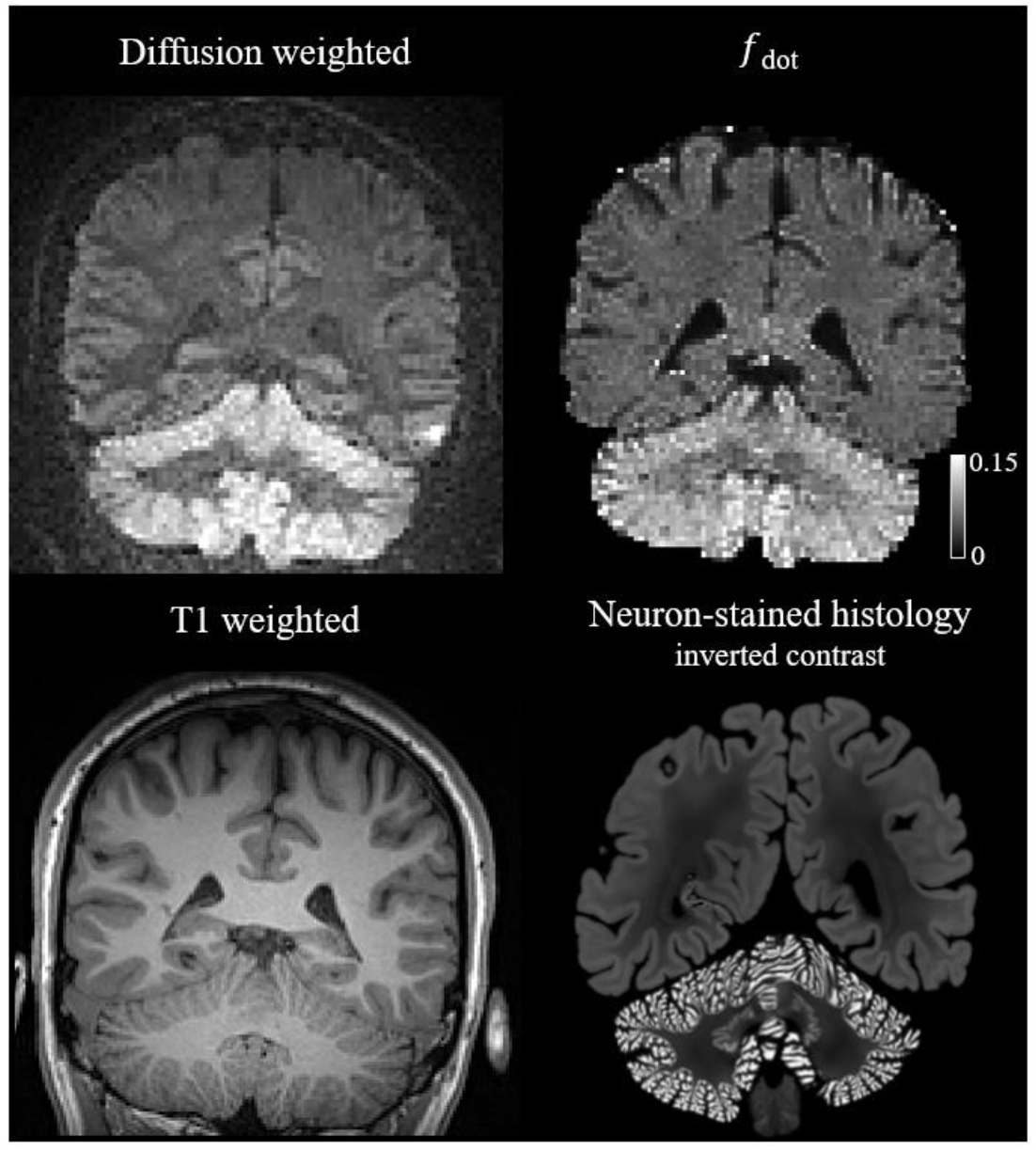
Signal retention using diffusion-weighted imaging with spherical encoding at *b* = 4 ms/μm^2^ (upper left) and estimation of *f*_dot_ (upper right) show agreement with neuron-stained histology (lower right plot shows human brain histology from the BigBrain atlas [44]). As expected, regions of high signal correspond to the cerebellar cortex where granule cells are densely packed, whereas the white matter is suppressed by the spherical diffusion encoding (lower left for morphological reference).

## Discussion

In this work, we investigated the value of super-resolution reconstruction and how it impacts signal accuracy and precision in diffusion MRI. We found that SRR produced increased accuracy in a challenging application that would not be feasible without it (Fig. 7). We also expect that the increase in signal accuracy by SRR will lead to an improved accuracy of diffusion parameters [15].

SRR improves the accuracy of the diffusion weighted signal by suppressing the rectified noise floor. The gain follows the law of diminishing returns—going from an aspect factor of 1 to 2 has a larger positive effect than going from 5 to 6 (Fig. 2). The result is noteworthy for two reasons. First, this gain is achieved without changing the pulse sequence, in contrast to methods relying on averaging before magnitude reconstruction [10]. Secondly, it is robust. This differs from postprocessing methods that have been proposed to remove the bias [47]–[50], which rely on prior knowledge of the data distribution. This can be challenging in vivo due to subject motion and eddy current distortions. However, unlike these methods, SRR does not completely remove the bias and the accuracy gain depends on factors like aspect factor and diffusivity. Nonetheless SRR is robust as signal bias is mostly avoided rather than corrected.

Signal precision can be preserved or improved using SRR, depending on the aspect factor, T1-weighting, and regularization. SNR efficiency increased for large aspect factors, however, effects of T1-relaxation become dominant at sufficiently high aspect factors (Fig. 4). Previous studies suggested larger aspect factor would lead to larger improvements in SNR efficiency [14], [19]. However, this was under the assumption of complete T1-recovery, which does not hold when relatively short repetition times are used. Generally, we see that a precision increase is limited and heavily dependent on acquisition, reconstruction, and intrinsic tissue parameters. To the best of our knowledge, no other studies have investigated precision in SRR at matched effective resolutions, something that arguably is needed for a fair comparison.

As the gain in precision from SSR was only modest, the major benefit of SRR is seen for low-SNR scenarios where accuracy can be substantially improved. For example, SRR can facilitate high resolution imaging and/or the use of strong diffusion encoding which is otherwise prohibited by noise floor effects. We demonstrated this by enabling high-resolution imaging at ultra-high b-values with spherical tensor encoding for the purposes of dot fraction imaging, which was not possible with direct sampling (Fig. 9). Dot fraction imaging visualizes densely packed cells located in the cerebellar cortex, a novel contrast that a recent study has showed in MRI systems with ultra-strong gradients and acquisition at a poor spatial resolution [6]. We believe that our method can help to study neurodegenerative diseases affecting these cells in a higher resolution than has been possible before. Current acquisition times are just below 10 minutes and can be further optimized to comply with clinical routine.

In high-SNR situations, SRR has few benefits compared with direct sampling. One remaining benefit is as a tool for acceleration when used in combination with a diffusion model as previously shown by van Steenkiste et al. [14] and Jeurissen et al. [15]. Here, the diffusion directions are subsampled over the acquired low resolution images, such that directional information needed to fit the model can be acquired in less time.

We identified some limitations of the present study, regarding the generality of our findings. First, we assume that a single point spread function with a certain FWHM describes the effective resolution of a given sample/reconstruction matrix, to enable resolution-matching of images. These profiles are location-dependent, and their shape differs among sampling matrices, possibly making comparisons between images captured in the SNR efficiency inaccurate. Second, we used the identity matrix as regularization matrix. Using image-dependent regularization can have benefits, such as edge-preservation [51]. In addition, our analysis does not include motion and eddy current correction in the SRR model, while perfect registration in SRR is of importance to obtain non-blurry high-resolution results. This has only potential consequences for our in vivo results, and could be addressed in future work. Third, the current analysis is done using T1-relaxation times at 3T, and will therefore differ somewhat for imaging systems with different field strengths [37]. As T1 generally scales with field strength, we expect less precision benefits at higher field strengths and vice versa.

In conclusion, we have presented a comprehensive analysis of SRR that outlined the major features influencing the precision and accuracy of the diffusion-weighted signal. We showcased the use of SRR in an extraordinarily challenging combination of high resolution and spherical tensor encoding with ultrahigh b-values, where SRR can suppress noise floor effects and recover high signal accuracy. We expect that the open-source tools developed herein will support future experimental design, such that both acquisition and reconstruction parameters can be optimized for specialized purposes.

## Acknowledgements

We thank Ben Jeurissen and Lipeng Ning for stimulating discussions. We thank Siemens Healthcare (Erlangen, Germany) for access to the pulse sequence programming environment. This research was financially supported by VR (2016-03443, 2020-04549), eSSENCE, Cancerfonden and The Swedish Prostate Cancer Federation.

## Appendix

### A. Derivation of SNR efficiency

We derive ρ as given in Eq. 15. We start from Eq. 13, stating

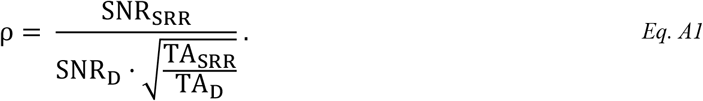

To find our final expression, we express both SNR_SRR_ and SNR_D_ in terms of SNR_LR_, the SNR of a low-resolution acquisition. For all cases,

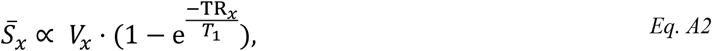

where 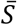 is the mean signal, *V* is the voxel size and *x* the reflecting component, i.e. LR, SRR or D (direct sampling). As *V*_LR_ = α · *V*_D_, we rewrite Eq. A2 to

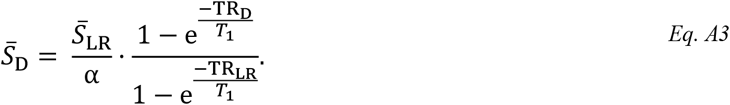

As σ is independent of voxel size, σ_d_ = σ_LR_ [27], combining both Eq. A3 and Eq. 12 gives

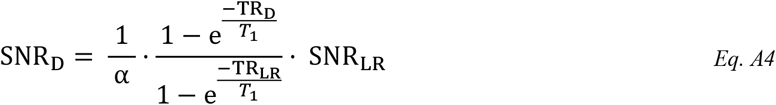

Along the same way we can find an expression for SNR_SRR_. As TR_SRR_ = TR_LR_, T1-effects are the same for both measurements, and we use Eq. A2 to see that

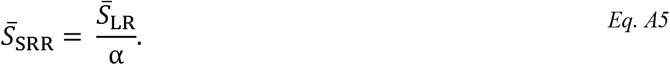

We use the definition of the noise propagation factor κ of Eq. 14 and Eq. 12 to find

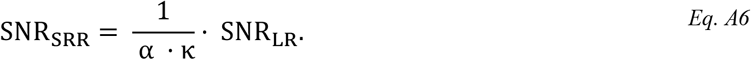

As for SRR the repetition time decreases, but the number of slice orientations increases with a factor *N_R_* we find

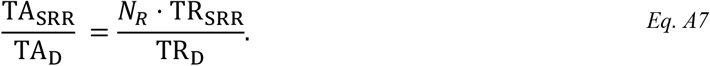

Combing Eq. A4, Eq. A6 and Eq. A7 in Eq. A1, gives the SNR efficiency ρ defined in Eq. 15, being

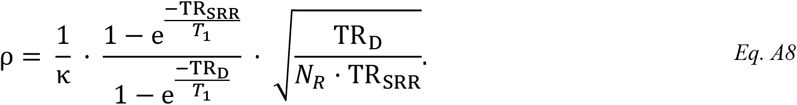

#### B. Gradient waveforms

Conventional, or linear diffusion encoding yields a pair of trapezoidal pulsed field gradient on each side of the refocusing pulse in a spin-echo sequence [1]. In this work, we used more advanced, spherical encoding where all three gradients are continuously used [34]. The gradient waveforms we used are depicted in Figure A1.

**Figure A1.**
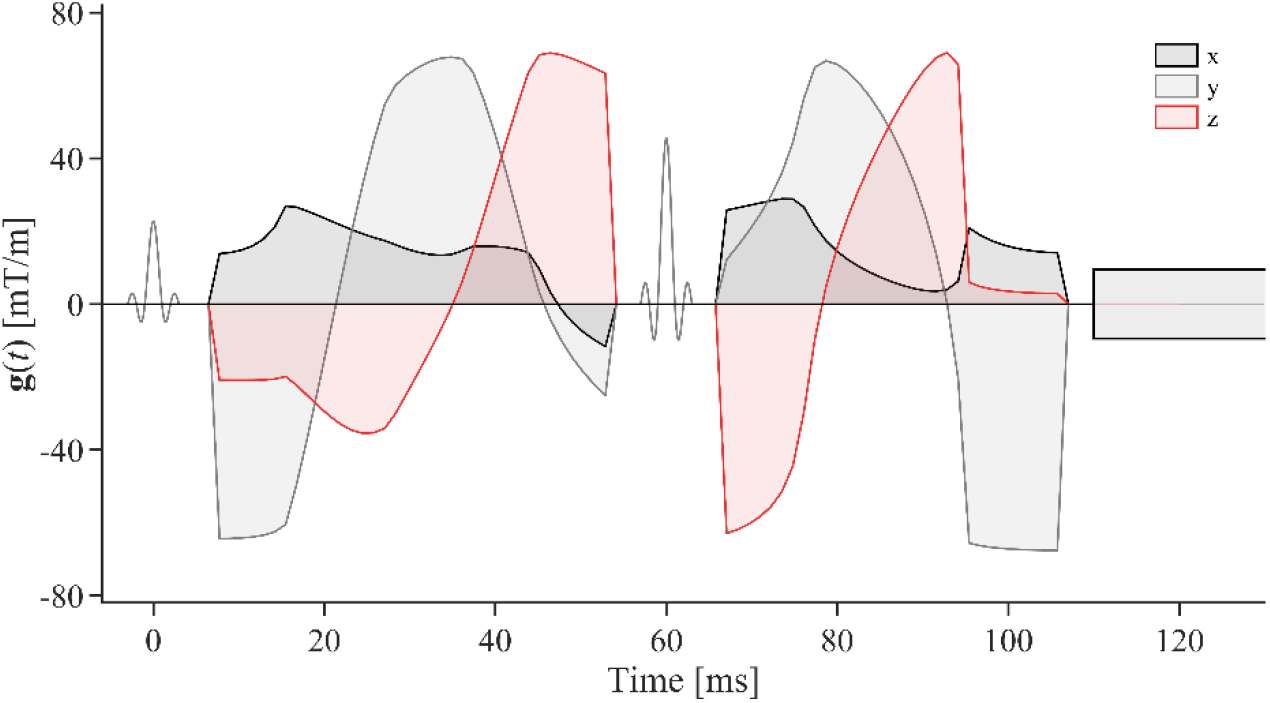
Optimized gradient waveforms were used to yield spherical tensor encoding (STE) in all practical experiments, and the one used for in vivo measurements is shown under conditions that yield *b* = 4.0 ms/μm^2^. Radiofrequency pulses (RF) are depicted in between gradients. This waveform and other resources related to the free waveforms sequence are available at https://github.com/filip-szczepankiewicz/fwf_seq_resources.

## Footnotes

1 We use the definition where accuracy is only a description of systematic errors. ISO calls this trueness [52].

